# Neural hyperactivity is a core pathophysiological change induced by deletion of an autism risk gene *Ash1l* in the mouse brain

**DOI:** 10.1101/2022.01.19.476965

**Authors:** Yuen Gao, Mohammad B Aljazi, Jin He

## Abstract

Autism spectrum disorder (ASD) is a neurodevelopmental disease associated with various gene mutations. Previous genetic and clinical studies report that mutations of the epigenetic gene *ASH1L* are highly associated with human ASD and intellectual disability (ID). Recent studies demonstrate that loss of *Ash1l* in the mouse brain is sufficient to induce ASD/ID-like behavioral and cognitive memory deficits, suggesting that disruptive *ASH1L* mutations are likely to be the causative driver leading to the ASD/ID pathogenesis in human patients. However, the brain pathophysiological changes underlying the *Ash1l*-deletion-induced ASD/ID-like behavioral and memory deficits remain unknown. Here we show loss of *Ash1l* in the mouse brain causes locomotor hyperactivity and higher metabolic rates . In addition, the mutant mice display lower thresholds for the convulsant reagent-induced epilepsy and increased neuronal activities in broad brain areas. Thus, our current study reveals that neural hyperactivity is a core pathophysiological change in the *Ash1l*-deficient mouse brain, which provides a brain-level basis for further studying the cellular and molecular mechanisms underlying the *Ash1l*-deletion-induced ASD/ID pathogenesis.

## Introduction

Autism spectrum disorder is one of most prevalent neurodevelopmental disorders that have a strong genetic basis(Lord et al., 2020). Recent genetic studies identified hundreds of ASD risk genes that are enriched with gene functions in epigenetically transcriptional regulation and chromatin modifications(Stessman et al., 2017; Grove et al., 2019; Ruzzo et al., 2019; Lalli et al., 2020; Satterstrom et al., 2020). Among the ASD risk genes encoding epigenetic factors, *ASH1L* (<u>*A*</u>bsent, <u>*S*</u>mall, or <u>*H*</u>omeotic discs <u>*1*</u>-<u>*L*</u>ike) is identified by multiple studies as a high ASD risk gene(Stessman et al., 2017; Grove et al., 2019; Ruzzo et al., 2019; Satterstrom et al., 2020). The genetic findings are further supported by multiple clinical cases reporting that some children diagnosed with ASD and/or ID acquire *de novo* disruptive or missense mutations of *ASH1L (*de Ligt et al., 2012; Wang et al., 2016; Okamoto et al., 2017; Faundes et al., 2018; Shen et al., 2019; Xi et al., 2020). In addition to ASD and ID, patients also display a variety of developmental abnormalities(de Ligt et al., 2012; Okamoto et al., 2017; Shen et al., 2019), suggesting critical roles of *ASH1L* in normal embryonic and postnatal development .

To examine the pathogenic role of disruptive *ASH1L* mutations in the ASD/ID genesis, a recent study used a conditional *Ash1l* knockout mouse line to show that deletion of *Ash1l* in the developing mouse brain was sufficient to cause multiple developmental defects, core autistic-like behaviors, and ID-like cognitive memory deficits, suggesting that the disruptive *ASH1L* mutations were likely to be the causative drivers leading to the human ASD/ID development(Gao et al., 2021). However, the pathophysiological changes in the *Ash1l-*deficient brain, which are critical for further understanding the molecular and cellular mechanisms underlying the *Ash1l*-deletion-induced ASD/ID pathogenesis, remain largely unknown.

In this study, we used the *Ash1l* brain-specific knockout mice to show that loss of *Ash1l* in the mouse brain causes locomotor hyperactivity, low body weight and reduced white adipose tissue depots with metabolic hyperactivity. Moreover, the mutant mice displayed lower thresholds for the convulsant reagent-induced epilepsy and increased neuronal activities in broad brain regions. These results suggest that neural hyperactivity is a core pathophysiological change in the *Ash1l*-deficient brain, which could play a critical role in leading to the brain functional abnormalities and ASD/ID-like behavioral and memory deficits.

## Materials and Methods

### Mice

The *Ash1l* conditional knockout mice were described in a previous report(Gao et al., 2021). Mice were housed under standard conditions (12 h light: 12 h dark cycles) with food and water ad libitum. All mouse experiments were performed with the approval of University Institutional Animal Care & Use Committee.

### Mouse breeding strategy

Generating *Ash1l*-Nestin-cKO mice: The *Ash1l* neural conditional knockout mice were generated by mating *Ash1l* floxed mice with Nestin-cre mice (B6.Cg-Tg (Nes-cre) 1Kln/J, The Jackson Laboratory). The wild-type (*Ash1l*^2f/2f^;Nestin-Cre^−/−^), heterozygous (*Ash1l*^2f/+^;Nestin-Cre^+/−^), and homozygous *Ash1l*-Nes-cKO (*Ash1l*-Nes-cKO, *Ash1l*^2f/2f^;Nestin-Cre^+/−^) were generated by *Ash1l*^2f/2f^;Nestin-Cre^−/−^ (female) x *Ash1l*^2f/+^;Nestin-Cre^+/−^ (male) mating.

### Genotyping

Genomic DNA was extracted from mouse tails with a lysis buffer of 0.01M NaOH. After neutralizing with Tris-HCl (PH 7.6), the extracted genomic DNA was used for genotyping PCR assays. Primers used for genotyping are the same as a previous report(Gao et al., 2021).

### Histology

Mouse adipose and liver tissues were fixed in 4% PFA in PBS at 4°C overnight and embedded in paraffin. For histology analysis, sections of 5 μm were stained with hematoxylin and eosin after dewaxing and rehydration. Images were captured using a Zeiss Axio Imager microscope (Carl Zeiss GmbH, Oberkochen, Germany) and an AxioCam HRc camera (Carl Zeiss GmbH). Adipocyte diameter and lipid droplet diameter were measured via Zeiss Zen Pro software (v.2.3; Carl Zeiss GmbH).

### Metabolic and activity analysis

Metabolic phenotyping and home cage activity were measured in TSE cages (PhenoMaster, TSE Systems) in Metabolic Core of Author University. Metabolic cage measurement is conducted continuously for 72 hours to account for acclimation of mice housed in home cages during this study. Mice were continuously monitored for food and water intake, locomotor activity, and energy expenditure. Ambient temperature was maintained at 20–23°C, and the airflow rate through the chambers was adjusted to maintain an oxygen differential around 0.3% at resting conditions. Metabolic parameters including VO2, respiratory exchange ratio, and energy expenditure were assessed via indirect calorimetry by comparing O2 and CO2 concentrations relative to a reference cage.

### Immunohistology assays

Mouse tissues were fixed in 4% PFA in PBS overnight at 4 °C and embedded in paraffin. For immunofluorescence, tissue sections of 5 μm were cut, dewaxed, and rehydrated. Antigen retrieval was performed by microwaving the sections in 0.01 M sodium citrate buffer (pH 6.0) for 4 min. Tissue sections were blocked in 5% normal donkey serum (NDS) for 30 min after washing with PBS. Tissue sections were then incubated with goat anti-Fos (1:300, sc-52-G, Cell Signaling technology) diluted in 5% NDS overnight at 4 °C. After washing with PBS, sections were incubated with Rhodamine (TRITC) AffiniPure donkey anti-goat IgG (1:300, Jackson ImmunoResearch Laboratories) for 1 h and mounted using Vectorshield mounting media with DAPI (H1200, Vector Laboratories). Images were captured using a Zeiss Axio Imager microscope (Carl Zeiss GmbH, Oberkochen, Germany) and an AxioCam HRc camera (Carl Zeiss GmbH) with image acquisition via Zeiss Zen Pro software (v.2.3; Carl Zeiss GmbH).

### EEG recording

2-month-old mice were used for the EEG recording, n = 3 males and 3 females for each genotype. Each animal was implanted with 3 recording electrodes placed at +1.50 A/P, +1.50 L; +1.50 A/P, −1.50 L; −2.0 A/P. +3.00 L, one ground electrode placed at −6.00 A/P, +0.50 L and one reference electrode placed at −3.5 A/P, +3.50 L relative to bregma. 14 days after operation, EEG was recorded on freely moving mice for 30 min before and after the PTZ (40mg/kg, i.p.) injection. The EEG data was recorded in the Neurologger2A. The EEG was analyzed by EEGLAB and visualized by EDFbrower version 1.88.

### Statistical analysis

All statistical analyses were performed using GraphPad Prism 8 (GraphPad Software). Parametric data were analyzed by a two-tailed *t* test or two-way ANOVA test for comparisons of multiple samples. *P* values < 0.05 were considered statistically significant. Planned comparisons (Šídák’s multiple comparisons test) were used if ANOVAS showed significant main or interaction effects. Data are presented as mean ± SEM.

## Results

### Loss of *Ash1L* in the mouse brain causes locomotor hyperactivity

To examine the causality between disruptive *ASH1L* mutations and autism pathogenesis, a recent study generated an *Ash1l* conditional knockout (*Ash1l*-cKO) mouse line and deleted *Ash1l* in the mouse brain by crossing the *Ash1l*-cKO mice with Nestin-Cre mice (*Ash1l*-Nes-cKO)(Gao et al., 2021). The Cre recombinase expressed in the Nestin+ neural progenitor cells (NPCs) induced *Ash1l* deletion in NPCs and NPC-derived neuronal and glial lineages, thus leading to *Ash1l* loss in the mouse brain. Using this mouse model, a recent study reported that loss of *Ash1l* in the mouse brain was sufficient to induce ASD-like social behavioral deficits, repetitive behaviors, and ID-like cognitive memory deficits, suggesting that disruptive *ASH1L* mutations were likely to be a causative driver leading to autism pathogenesis in human patients. Notably, it was observed that the mutant mice had high locomotor activities compared to their wild-type littermates in the open field tests(Gao et al., 2021). To confirm this observation, we used the TSE PhenoMaster/LabMaster System to measure locomotor activities by counting total times that mice passed through an infrared sensor and total running distances in the cage(Van Klinken et al., 2012). The results showed that except for the male in the light-off cycle (ZT12-ZT0, Zeitgeber Time), both male and female mutant mice had significant more times passing through the infrared sensor (male, F(1,52) = 29.56, light cycle *p* < 0.001, dark cycle *p* = 0.319; female, F(1,52) = 259.0, light cycle *p* <0.001, dark cycle *p* < 0.001) and longer running distances (male, F(1,52) = 27.45, light cycle *p* < 0.001, dark cycle *p*= 0.1112; female, F(1,52) = 257.6, light cycle *p* < 0.001, dark cycle *p* < 0.001) (Fig. 1), which confirmed the previous observation that loss of *Ash1l* in the mouse brain caused higher locomotor activities(Gao et al., 2021). Notably, compared to wild-type controls, both male and female mutant mice had higher locomotor activities during the light-on cycle (ZT0-ZT12) in which wild-type mice spent most time in sleeping and maintained relatively low locomotor activities, suggesting the mutant mice had high locomotor activity during normal sleeping time, which was consistent with the sleeping difficulty commonly observed in human ASD patients.

**Figure 1.**
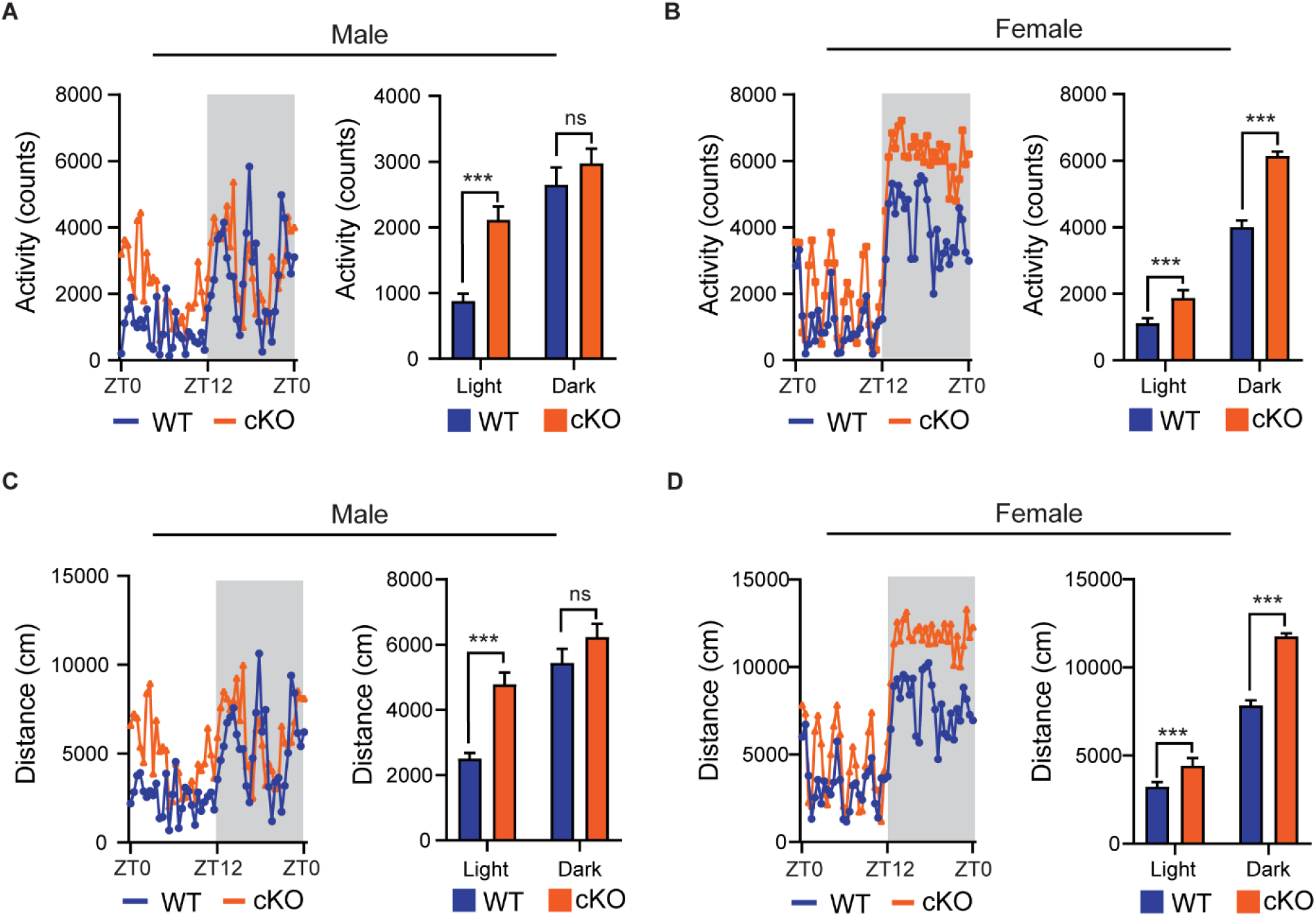
Loss of *Ash1L* in the mouse brain causes locomotor hyperactivity. **(A-B)** Plots showing total times (counts) that male (A) and female (B) wild-type and *Ash1l*-Nes-cKO mice passing through an infrared sensor in a 12-hour light-on cycle (ZT0 – ZT12, Zeitgeber Time) and a 12-hour light-off cycle (ZT12 – ZT0, Zeitgeber Time). n = 4 males and 4 females per genotype. *P*-values calculated using two-way ANOVA test. Error bars in graphs represent mean ± SEM. \*\*\**p* < 0.001; ns, not significant. **(C-D)** Plots showing the distances that male (C) and female (D) wild-type and *Ash1l*-Nes-cKO mice run in the 12-hour light-on cycle (ZT0 – ZT12, Zeitgeber Time) and the 12-hour light-off cycle. n = 4 males and 4 females per genotype. *P*-values calculated using two-way ANOVA test. Error bars in graphs represent mean ± SEM. Note: \*\*\**p* < 0.001; ns, not significant.

### Loss of *Ash1l* in the mouse brain reduces adipose tissue depots

In addition to the postnatal growth retardation observed in the *Ash1l* mutant pups(Gao et al., 2021), the adult mutant mice had significantly lower body weight compared to their wild-type littermates (t = 4.688, df = 7, *p* = 0.0022) (Fig.2a,b). Consistently, the mutant mice had markedly reduced subcutaneous (t = 7.259, df = 7, *p* = 0.0002) and visceral adipose tissue depots (t = 7.180, df = 7, *p* = 0.0002), which appeared to mainly affect white adipose tissues (WATs), a major form of adipose tissues for triacylglycerol storage(Saely et al., 2012), but not brown adipose tissues (BATs) (t = 0.0150, df = 7, *p* = 0.9884) (Fig.2c-g). Histological analyses showed that the WATs in the mutant mice had smaller cell sizes (t = 11.49, df = 58, *p* < 0.0001) but maintained the cell numbers (Fig.2h, i), suggesting that reduced WATs in the mutant mice were likely to be caused by energy over-expenditure and reduced triacylglycerol storage in WATs but not due to the loss of adipocytes.

**Figure 2.**
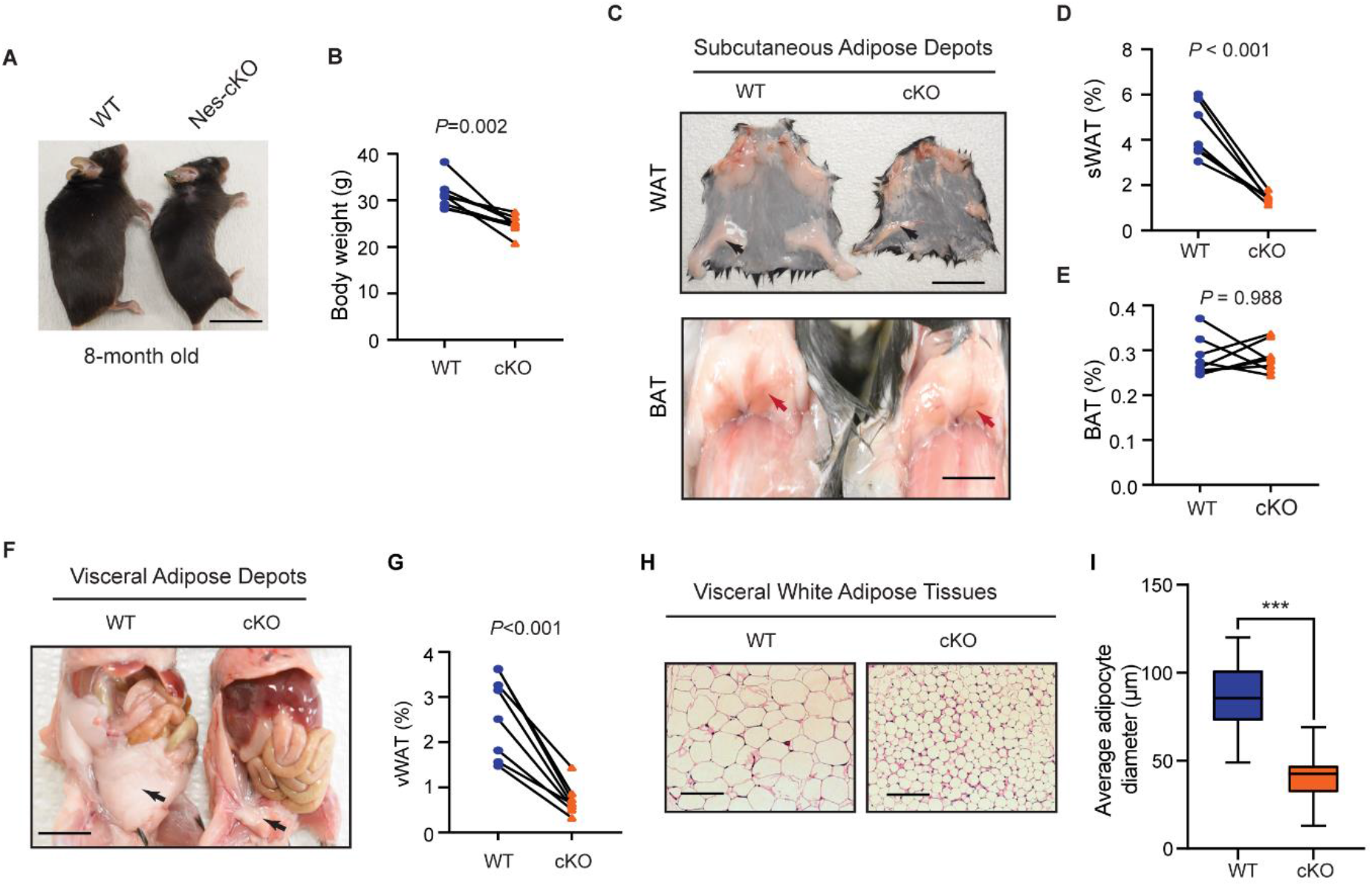
Loss of *Ash1l* in the mouse brain reduces adipose tissue depots. **(A)** Representative photos showing the low body weight of adult *Ash1l*-Nes-cKO mice compared to wild-type littermates. (**B**) Plot showing the body weight of adult (8-month-old) wild-type and *Ash1l*-Nes-cKO mice. For each group, n = 8. *P*-values calculated using paired *t* test. (**C**) Photos showing subcutaneous white adipose depots (WAT) and brown adipose deports (BAT) in wild-type and *Ash1l*-Nes-cKO adult mice. bar = 0.5 cm. (**D-E**) Plots showing the quantitative measurement of subcutaneous white adipose depots (sWAT) and brown adipose depots (BAT) in wild-type and *Ash1l*-Nes-cKO mice. Y-axis represents the percentage of subcutaneous adipose tissue normalized to total body weight. For each group, n = 8. *P*-values calculated using paired *t* test. (**F**) Representative photos and showing the visceral white adipose depots in wild-type and *Ash1l*-Nes-cKO mice. bar = 0.5 cm. (**G**) Plots showing the quantitative measurement of visceral white adipose depots (vWAT) in wild-type and *Ash1l*-Nes-cKO mice. Y-axis represents the percentage of visceral adipose tissue normalized to total body weight. For each group, n = 8. *P*-values calculated using paired *t* test. (**H**) Photos showing the adipocytes of visceral white adipose tissues under microscope. Bar = 50 μM. (**I**) Plot showing the average diameters of adipocytes of wild-type and *Ash1l*-Nes-cKO mice. For each group, n= 30. *P*-values calculated using unpaired *t* test.

### Loss of *Ash1l* in the mouse brain causes metabolic hyperactivity

To examine the underlying reasons leading to the reduced white adipose tissue depots in adult mutant mice, we used the TSE PhenoMaster/LabMaster System to examine overall metabolism by measuring the calorimetry components including food intake, oxygen (VO_2_) consumption, carbon dioxide (VCO_2_) production, respiration exchange ratio (VCO_2_/VO_2_), and heat generation. The results showed that both male and female mutant mice had higher or similar food intake compared to their wild-type littermates (male, F(1,12) = 13.49, light cycle *p*= 0.0322, dark cycle *p*= 0.1150; female, F(1,12) = 25.08, light cycle *p* = 0.7243, dark cycle *p* = 0.0010) (Fig.3a, b), suggesting the reduced WAT depots were not caused by insufficient food intake. Consistent with the high locomotor activities, the mutant mice had higher oxygen consumption (F(1,52) = 40.19, light cycle *p* < 0.0001, dark cycle *p* <0.0001), carbon dioxide production (F(1,52) = 43.54, light cycle *p* < 0.0001, dark cycle *p* < 0.0001), respiration exchange ratios (F(1,52) = 48.00, light cycle *p* < 0.0001, dark cycle *p* = 0.0058), and heat generation (F(1,52) = 41.58, light cycle *p* < 0.0001, dark cycle *p* < 0.0001) (Fig.3c-f), suggesting loss of *Ash1l* in the brain was likely to be caused overall metabolic hyperactivity, which led to a high calorie consumption and lipid catabolism in adult mutant mice. Notably, the metabolic hyperactivity of mutant mice appeared to be more prominent during the light-on cycle (ZT0-ZT12) in which wild-type mice spent most time in sleeping and had a relatively low metabolic rate, further suggesting that the normal circadian cycle and sleep were disturbed in the mutant mice.

**Figure 3.**
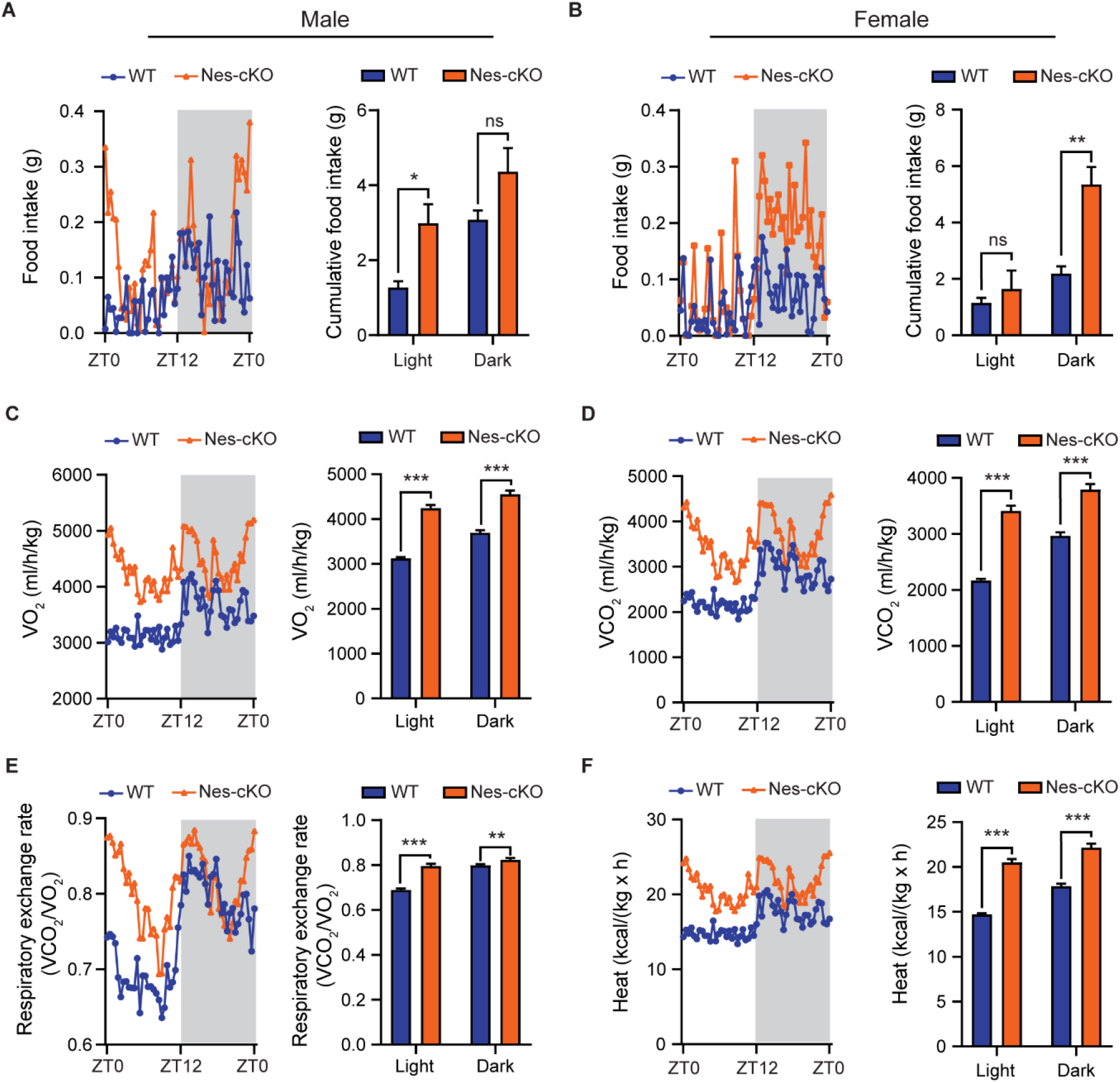
Loss of *Ash1l* in the mouse brain causes metabolic hyperactivity. **(A-B)** Plots showing the food intake of male (A) and female (B) wild-type and *Ash1l*-Nes-cKO mice in the light-on cycle (ZT0 – ZT12, Zeitgeber Time) and in light-off cycle (ZT12 – ZT0). n = 4 males and 4 females per genotype. *P*-values calculated using two-way ANOVA test. Error bars in graphs represent mean ± SEM. \**p* < 0.05; \*\**p* < 0.01; ns, not significant. **(C-F)** Plots showing the oxygen (VO2) consumption (C), carbon dioxide (VCO2) production (D), respiration exchange ratio (VCO2/VO2) (E), and heat generation (F) of wild-type and *Ash1l*-Nes-cKO mice in the 12-hour light-on cycle (ZT0 – ZT12, Zeitgeber Time) and in the 12-hour light-off cycle (ZT12 – ZT0, Zeitgeber Time). n = 4 males and 4 females per genotype. For each group, n = 27. *P*-values calculated using two-way ANOVA test. Error bars in graphs represent mean ± SEM. \*\**p* < 0.01; \*\*\**p* < 0.001.

### Loss of *Ash1l* in the mouse brain reduces the thresholds for the pentylenetetrazole (PTZ)-induced epilepsy

Previous studies have shown that ASD patients have a high occurrence of epilepsy(Besag, 2018; Pacheva et al., 2019). Similarly, epilepsy is also observed in the patients with *ASH1L* mutations(de Ligt et al., 2012; Shen et al., 2019). Although the mutant mice did not have spontaneous epilepsy, the locomotor and metabolic hyperactivity observed in the mutant mice suggested that loss of *Ash1l* in the brain had reduced inhibitory signals and shifted the excitation/inhibition balance towards over-excitation in cortices, which could lower the thresholds for convulsant reagent-induced epilepsy. To test this hypothesis, we challenged both wild-type and mutant mice by intraperitoneal injection of PTZ, a GABAA receptor antagonist, at a sub-convulsant dose (40mg/kg) to induce seizures(Sansig et al., 2001). The thresholds for the PTZ-induced seizures were measured by scoring epileptic behaviors in 10 minutes after PTZ administration according to the previous report(Van Erum et al., 2019). The results showed that the sub-convulsant dose of PTZ induced minor epileptic behaviors such as moving arresting, whisker trembling, and facial jerking in wild-type mice. In contrast, both male (t = 14, df = 10, *p* < 0.0001) and female ( t = 10.5, df = 10, *p* = 0.0001) mutant mice displayed much more severe epileptic behaviors including heavy myoclonic jerks, lying on belly with rapid body twitches, and clonic-tonic spasm, suggesting that the mutant mice had lower thresholds for the PTZ-induced epilepsy (Fig. 4a). Moreover, electroencephalography (EEG) recordings on frontal cortices showed that although both wild-type and mutant mice had reduced electric wave frequency after PTZ administration, the mutant mice had spike-wave electrical discharges with increased amplitude (Fig. 4b), which was consistent with the severe epileptic behaviors observed in the mutant mice. Altogether, the results suggested that loss of *Ash1l* in the mouse brain increased neural activities in cortices, which decreased the thresholds for the convulsant reagent-induced epilepsy.

**Figure 4.**
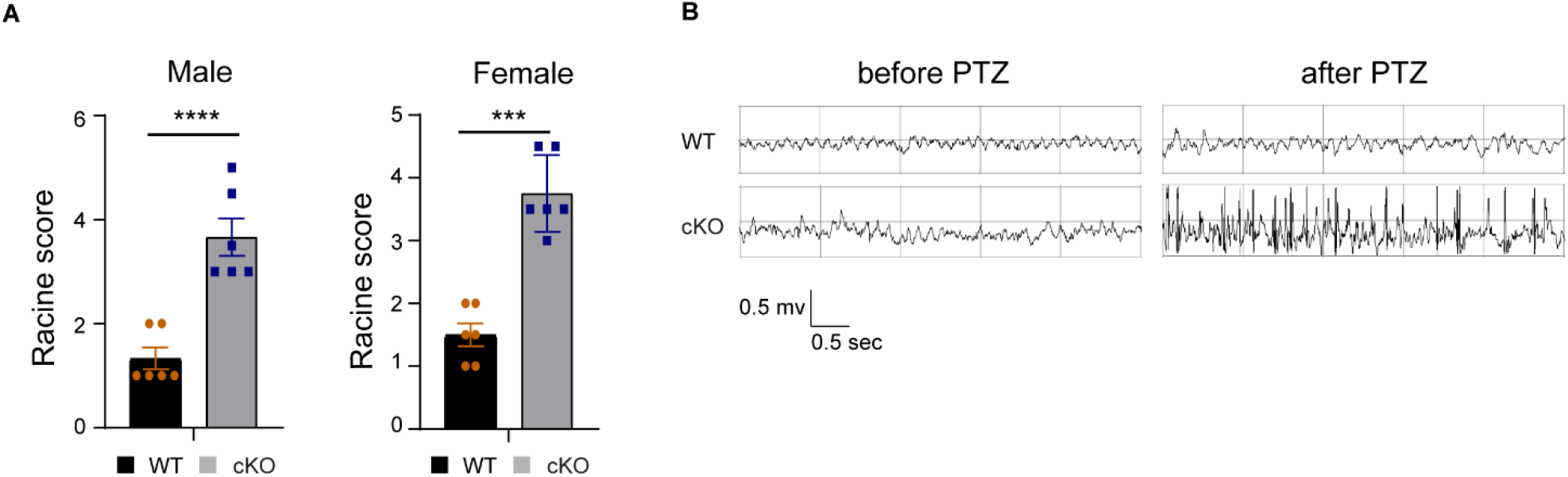
Loss of *Ash1l* in the mouse brain reduces the thresholds for the PTZ-induced epilepsy. **(A)** Plots showing the Racine score(Van Erum et al., 2019) of epileptic behaviors of male and female wild-type and *Ash1l*-Nes-cKO mice. n = 6 males and 6 females per genotype. *P* value calculated using a two-tailed *t* test. Error bars in graphs represent mean ± SEM. \*\*\**p* < 0.001. **(B)** Representative EEG recording showing the electrical waves before and after sub-convulsant dose of PTZ injection.

### Loss of *Ash1l* in the mouse brain causes neuronal hyperactivity in broad brain regions

To identify brain areas involved in locomotor and metabolic hyperactivity in the mutant mice, we used the c-Fos immunoreactivity as a neuronal activation marker to screen brain regions with neuronal hyperactivity(Bullitt, 1990). The results showed that compared to the wild-type controls, the mutant mice had significant increased c-Fos+ neurons in motor cortices (t = 6.1, df = 4, *p* = 0.0037), amygdala (t = 9.5, df = 4, *p* = 0.0007), and hypothalami (t = 4.8, df = 4, *p* = 0.0084) (Fig. 5a-f), consistent with locomotor hyperactivity, increased anxiety-like behaviors(Gao et al., 2021), and metabolic hyperactivity observed in the mutant mice.

**Figure 5.**
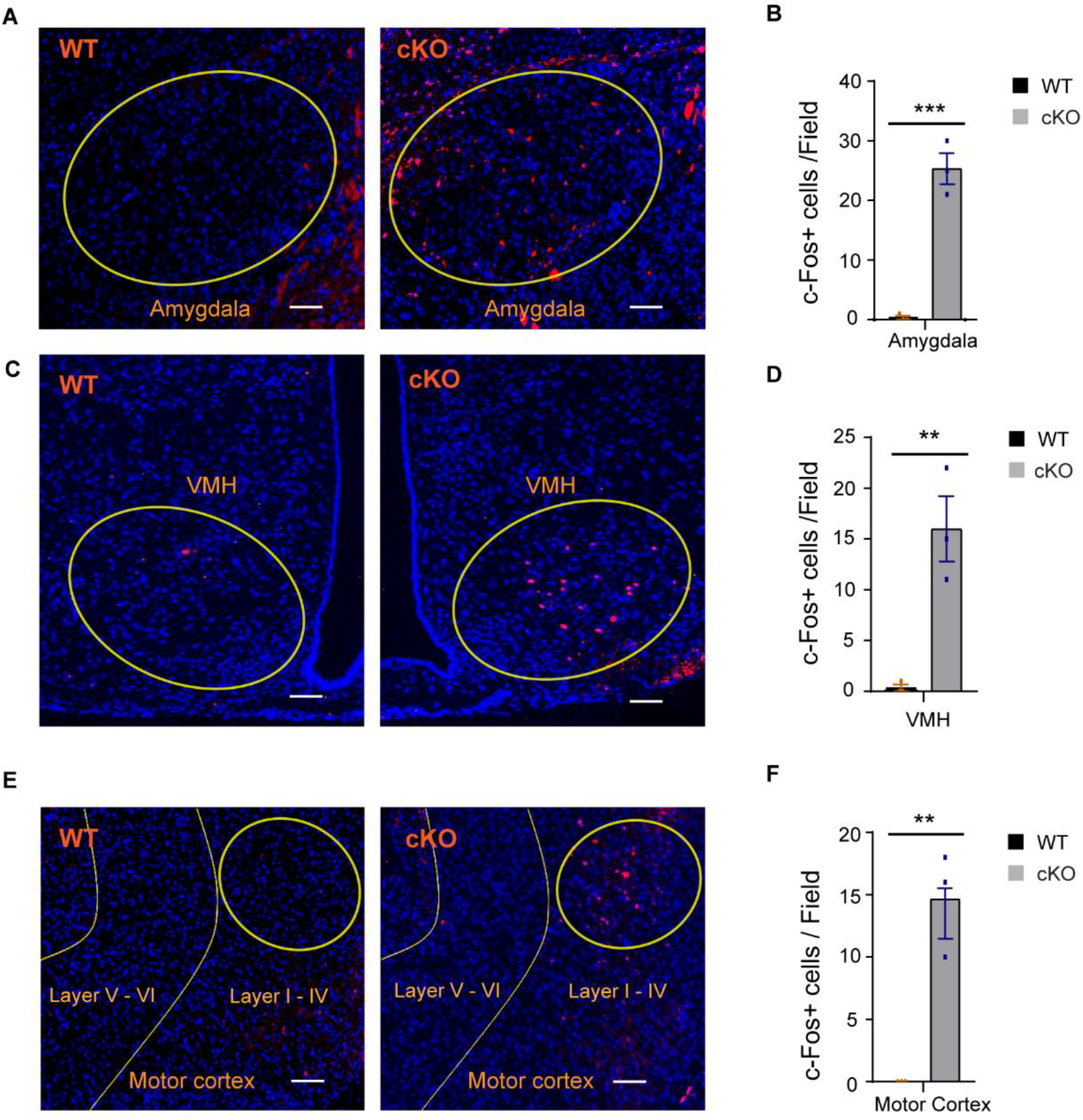
Loss of *Ash1l* in the mouse brain causes neuronal hyperactivity in multiple brain regions. **(A-F)** Photos (A, C, E) and bar plots (B, D, F) showing the c-Fos+ cells in amygdala (A, B), ventromedial hypothalamus (VMH) (C, D), and motor cortices (E, F). n = 3 per genotype. *P* value calculated using a two-tailed *t* test. Error bars in graphs represent mean ± SEM. \*\*\**p* < 0.001. bar = 100 μm.

## Discussion

ASH1L is a histone methyltransferase that mediates di-methylation of histone H3 lysine 36(Yuan et al., 2011). Functionally, ASH1L facilitates gene expression through its epigenetic modification of histone(Schuettengruber et al., 2011). Recent large-scale genetic studies reveal that *ASH1L* is a high ASD risk gene, which are supported by multiple clinical case reports that various *ASH1L* gene mutations are identified in children diagnosed with ASD/ID. To examine the causality between *ASH1L* mutations and ASD pathogenesis, a recent study used the *Ash1l* conditional knockout mice to show that loss of *Ash1l* in the mouse brain was sufficient to induced core ASD-like behavioral deficits, including impaired sociability, reduced social memory, repetitive behaviors, as well as ID-like cognitive memory deficits, suggesting disruptive *ASH1L* mutations were likely be the causative driver leading to ASD/ID genesis(Gao et al., 2021). However, the pathophysiological changes in the *Ash1l*-deficient brain that underline the autistic-like behavioral deficits remain unclear.

In this study, we provide multiple lines of evidence to show that neural hyperactivity is a core pathophysiological change in the *Ash1l*-deficient mouse brain. First, consistent with the observation that the *Ash1l* mutant mice had longer running distances in the open field test(Gao et al., 2021), the 72-hour locomotor activity test showed that compared to wild-type controls, the *Ash1l* mutant mice had increased movement in cages and longer running distances, confirming that loss of *Ash1l* in the mouse brain caused locomotor hyperactivity (Fig. 1). Second, the adult *Ash1l* mutant mice had lower body weight and markedly reduced white adipose tissue depots, which was accompanied with significantly reduced size of adipocytes (Fig. 2). The systemic measurement of metabolic activities demonstrated that both male and female *Ash1l* mutant mice had increased food intake and metabolic rates (Fig. 3), suggesting the lower body weight and reduced adipose tissues of *Ash1l* mutant mice were likely to be the secondary effect of hyperactivity-induced energy over-expenditure but not the hypothalamus-deficiency-induced insufficient food intake. Notably, the mutant mice maintained both locomotor and metabolic hyperactivities during the normal sleeping cycle, suggesting that normal sleep was disturbed in the *Ash1l* mutant mice due to neural hyperactivity (Figs. 1, 3), which was consistent with the sleep difficulty commonly observed in ASD patients(Devnani and Hegde, 2015; Elrod and Hood, 2015; Veatch et al., 2015). Third, although the *Ash1l* mutant mice did not develop spontaneous epilepsy, we observed that the mutant mice had reduced thresholds for the convulsant reagent-induced epilepsy. Compared to wild-type controls, the mutant mice had more severe epileptic behaviors and electrical discharges in cortices in response to a sub-convulsant dose of PTZ (Fig. 4). Finally, compared to wild-type controls, the *Ash1l* mutant mice had increased c-Fos+ neuronal populations in multiple brain regions (Fig. 5), suggesting the *Ash1l*-deletion-induced neuronal hyperactivity occurred in broad cortical and subcortical areas but not restricted to specific brain regions.

Increased ratio of excitatory versus inhibitory neural signals has been found to be a common change in ASD patients associated with a variety of genetic and environmental factors (Uzunova et al., 2016; Goel and Portera-Cailliau, 2019). It has been postulated that increased excitation/inhibition (E/I) signals generate a higher background of noise, which interferes the development and maturation of neural network in the developing brain(Rubenstein and Merzenich, 2003; Sohal and Rubenstein, 2019). Consistent with this theory, our current study demonstrates that neural hyperactivity is a core pathophysiological change in the brain with loss of *Ash1l*, a high ASD risk gene identified in human patients. Although the mechanisms underlying the *Ash1l*-mutation-related ASD behavioral deficits remain largely unelucidated, the identification of core brain pathophysiological changes in the *Ash1l*-deficient brain provides an organ-level basis for further dissecting the molecular and cellular components contributing to the E/I imbalance caused by *Ash1l* loss in the brain.

## Acknowledgements

We thank Drs. Alfred J. Robison and Gina Leinninger for experimental data interpretation and discussion. This work was supported by the National Institutes of Health (grant R01GM127431).

## Author Contributions

J.H. conceived the project. Y.G. and J.H. performed the experiments. Y.G. and M.B.A. maintained the mouse colonies. Y.G. and J.H. wrote the manuscript.

## Competing Interest Statement

Authors declare no competing interests.

## Notes

### Competing Interest Statement

The authors have declared no competing interest.

